# Adaptative Laboratory Evolution reveals biofilm regulating genes as key players in *B. subtilis* root colonization

**DOI:** 10.1101/2023.07.04.547689

**Authors:** Maude Pomerleau, Vincent Charron-Lamoureux, Lucille Léonard, Frédéric Grenier, Sébastien Rodrigue, Pascale B. Beauregard

## Abstract

Root-associated microorganisms play an important role in plant health, such as plant growth-promoting rhizobacteria from the *Bacillus* and *Pseudomonas* genera. Although bacterial consortia including these two genera would represent a promising avenue to efficient biofertilizer formulation, we observed that *B. subtilis* root colonization is decreased by the presence of *P. fluorescens* and *P. protegens*. To determine if *B. subtilis* can adapt to the inhibitory effect of *Pseudomonas* on roots, we conducted adaptative laboratory evolution experiments with *B. subtilis* in mono-association or co-cultured with *P. fluorescens* on tomato plant roots. Evolved isolates with various colony morphology and stronger colonization capacity of both tomato plant and *A. thaliana* roots emerged rapidly from the two evolution experiments. Certain evolved isolates had also a better fitness on root in presence of other Pseudomonas species. Whole genome sequencing revealed that single nucleotide polymorphism (SNPs) in negative biofilm regulator genes *ywcC* or *sinR* were found in all independent lineages, suggesting their involvement in enhanced root colonization. These findings provide insights into the molecular mechanisms underlying *B. subtilis* adaptation to root colonization and highlight the potential of directed evolution to enhance beneficial traits of PGPRs.

## Introduction

Numerous microorganisms live and thrive close to plant roots and form a distinct microbiota playing an important role in promoting plant health (1). Certain plants were shown to actively select beneficial bacteria in the rhizosphere via the secretion of specific compounds in their exudates (2), stimulating an interaction that benefit both the plants and bacteria. One of these microorganisms is *Bacillus subtilis*, a Gram-positive, soil ubiquitous bacteria that was define more than 40 years ago as a plant growth-promoting rhizobacteria (PGPR) (3). Many strains from the *Bacillus subtilis* species complex are widely used in agricultural settings due to their multiple beneficial effects on plants (4–7). They can have direct effects, such as increasing nutrient bioavailability and stimulating plant growth via the production of phytohormones (8,9). They can also serve as biocontrol agents, since secondary metabolites such as plipastatin and surfactin have antimicrobial activity against pathogens (7,10–12). *B. subtilis* can also prime the plant defense system by triggering the induced systemic resistance (2).

While many PGPR strains show strong, reproducible plant enhancements in controlled conditions, biofertilizers effects are variable in field conditions (13–16). Different factors can explain this lack of efficacy, such as the extent of the inoculant biological activity in the receiving matrices (17,18). For example, *Bacillus subtilis* was shown to rapidly sporulate when cells were introduced in bulk soil, thereby limiting their beneficial effect. Root exudates of certain plants, exposed or not to other microorganisms, can modulate this metabolically inactive state (19,20). A second factor is the inoculant capacity to establish in the rhizosphere in presence of the residing microbiota (17,18). Multiple interactions with the soil community can result in the inhibition of important traits for *B. subtilis* rhizosphere establishment (21,22).

One key factor for *B. subtilis* root colonization is its capacity to form biofilms on roots (6,23,24). The biofilm extracellular matrix of *B. subtilis* is composed of exopolysaccharides (EPS), protein fibers (TasA and TapA) and a small hydrophobin protein (BslA) (4,25). Poly-y-glutamic acids (y-PGA), a secreted polymer, was also reported to play an important role in biofilm robustness and root colonization (26). In non-biofilm conditions, operons encoding for EPS synthesis and TasA/TapA are repressed by the transcriptional repressor SinR (4,25). Plant polysaccharides and malic acids, which are respectively present at the surface of the root and in root exudates, trigger a signalling pathway resulting in SinR inhibition and expression of matrix genes (23,27).

Adaptative Laboratory Evolution (ALE) experiments are a powerful tool to study adaptations of microbes to an environment (28). Performed in mono-association with a plant, these assays can lead to the emergence of strong mutualists traits and reveal the molecular mechanisms involved in colonization (29–32). Laboratory evolution of *B. subtilis* on *A. thaliana* led to the emergence of evolved isolates of various morphologies with increased root colonization by themselves or in combination (30,31). Some mutations were found to influence biofilm formation in presence of xylan and decreased motility, suggesting an evolutionary trade-off between those traits (30,32). One isolate was also shown to increase root colonization within a synthetic community (30).

Many *Pseudomonas* species are also characterized as PGPR, and several were co-isolated with *Bacillu*s from the rhizosphere where they likely interact with each other (33–35). Even if the combination of these two genera can increase biocontrol abilities and plant growth, most interactions between them were reported to be antagonistic *in vitro* (22,36–39). For example, certain *Pseudomonas* can repress or disperse *B. subtilis* biofilm via the production of the secondary metabolite 2,4-diacetylphloroglucinol (DAPG) and rhamnolipids (21,40,41). These interactions likely impact *B. subtilis* capacity to strongly establish on plant roots.

In this study, we first observed that *Pseudomonas* species impaired the establishment of *B. subtilis* on the roots of *A. thaliana* and tomato seedlings. To determine if *B. subtilis* can evolve to overcome this challenge and what traits would be involved, we elaborated an ALE on tomato roots in which *B. subtilis* was challenged or not by *Pseudomonas fluorescens*. Evolved isolates with enhanced root colonization rapidly emerged. Multiple mutations were identified in the negative regulators of biofilm formation *ywcC* and *sinR* for both evolution experiments, which suggests that the ability to compete with *Pseudomonas* for root colonization is highly linked to biofilm formation.

## Results

### *Pseudomonas* spp. impede *B. subtilis* colonization on seedlings

*In vitro* antagonism between *Pseudomonas* species and *B. subtilis* was often reported, but no studies have yet shown how these interactions translated on plant roots (21,22,36,42). We thus performed root colonization assays with *B. subtilis* NCIB 3610 in presence of three *Pseudomonas* species that are used as biocontrol agents (*P. fluorescens* WCS365, *P. capeferrum* WCS358, and *P. protegens* Pf-5) (43,44). *Arabidopsis thaliana* and tomato, grown in hydroponic conditions, were used for this assay since the outcome might be plant specific. *B. subtilis* colonization was quantified by colony-forming unit (CFU) enumeration for *A. thaliana*, while the dense biofilm formed on tomato roots made this technique imprecise. Consequently, we used an alternative method where *B. subtilis* constitutively expressed the fluorescence gene *mKate2* and the relative fluorescence unit present on the root was measured. As shown in Fig 1, *P. protegens* and, in a lesser extent, *P. fluorescens*, decreased *B. subtilis* presence on both plants, while *P. capeferrum* influenced *B. subtilis* establishment on *A. thaliana* only. These results points toward a general, plant-independent inhibitory effect of *Pseudomonas* on *B. subtilis* root colonization.

**Fig 1.**
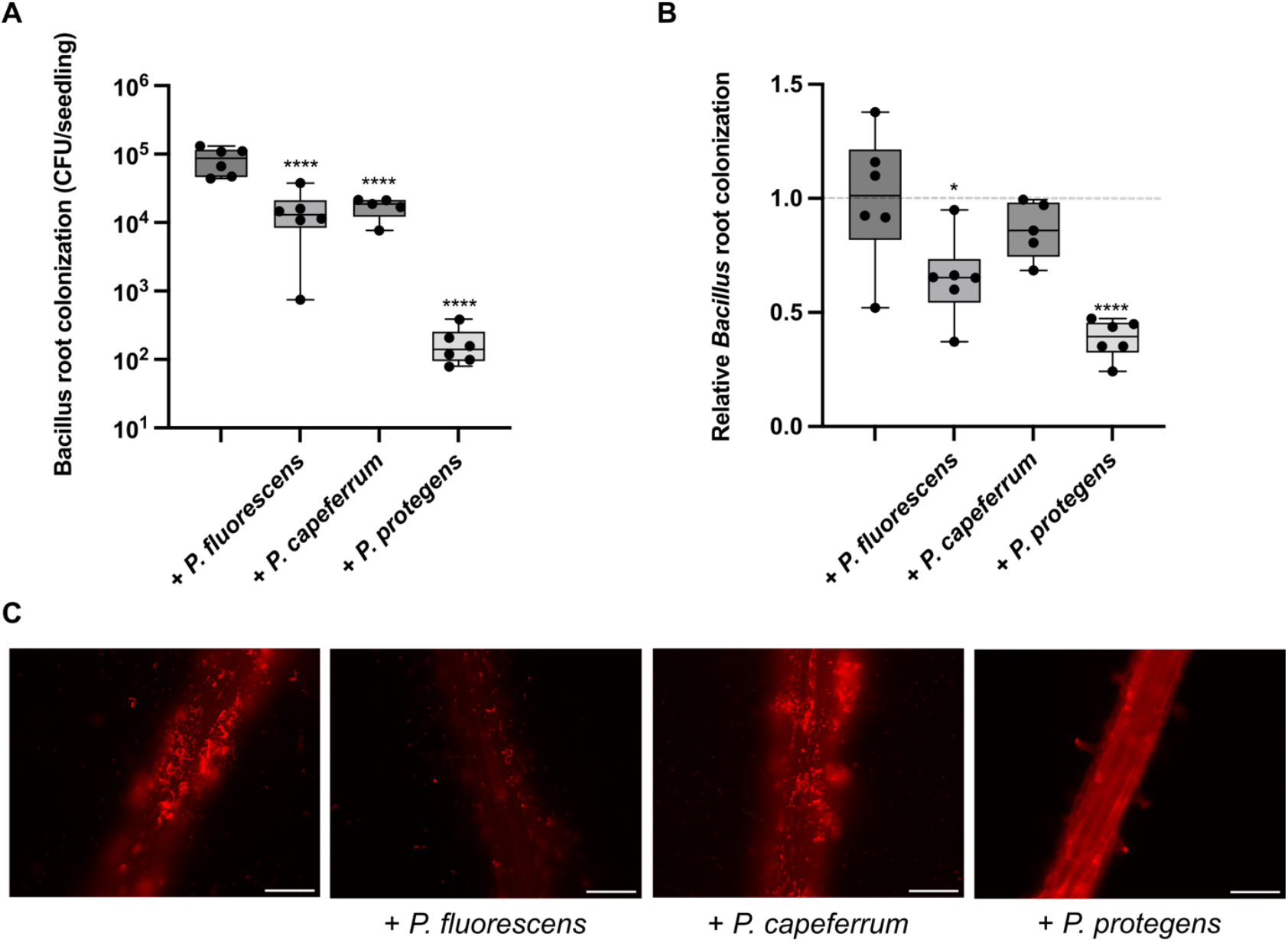
*B. subtilis* colonization is decreased by *Pseudomonas* species. (A) Quantification of *B. subtilis* colonization on 6 days-old *A. thaliana* roots when inoculated alone or with different *Pseudomonas*. (B) Relative quantification of tomato root colonization by *B. subtilis* NCIB 3610 constitutively expressing mKATE2 and inoculated with *Pseudomonas* spp., reported as a ratio over the mean fluorescence intensity of *B. subtilis* colonizing the plant alone * = P < 0.05, **** = P < 0.0001, ANOVA followed by Dunnett’s post hoc test. (C) Representative pictures of *A. thaliana* root colonization when *B. subtilis* is inoculated alone or with different *Pseudomonas*. Scale bar is 100 μm for all images.

### *Bacillus subtilis* evolves rapidly on tomato plant roots

To determine if *B. subtilis* could adapt to root colonization in presence of a competitor, we performed a directed evolution assay in which we selected for *B. subtilis* cells able to colonize the root. Briefly, the ancestral strain *B. subtilis* NCIB 3610 was inoculated on tomato roots before being transferred to a new root every 24 h for a total of 21 cycles (Fig 2A). The experiment was performed with *B. subtilis* alone (Bacillus-Root Evolution; BRE) or with the addition of *Pseudomonas fluorescens* WCS365 acting as a competitor (Bacillus-Root-Pseudomonas Evolution; BRPE). This strategy was chosen to help determine if the acquired mutations were specific for improved colonization, competition or shared between both evolution assays. *P. fluorescens* was chosen as the competitor strain in the BRPE experiment due to its moderate impact on *B. subtilis* colonization, as opposed to *P. protegens* which completely outcompeted *B. subtilis* during an evolution attempt (data not shown). We followed colonization of the six lineages (6 independent plants) at each cycle by evaluating the colonization of *B. subtilis* on root (Fig 2B and 2C). Of note, one of the lineages with *P. fluorescens* was lost due to contamination and is consequently not shown. Interestingly, the BRPE showed an upward trend in colonization capacity, while the BRE had a drop in the first cycles and ended approximately at the same colonization capacity than the starting point (Fig 2B and 2C).

**Fig 2.**
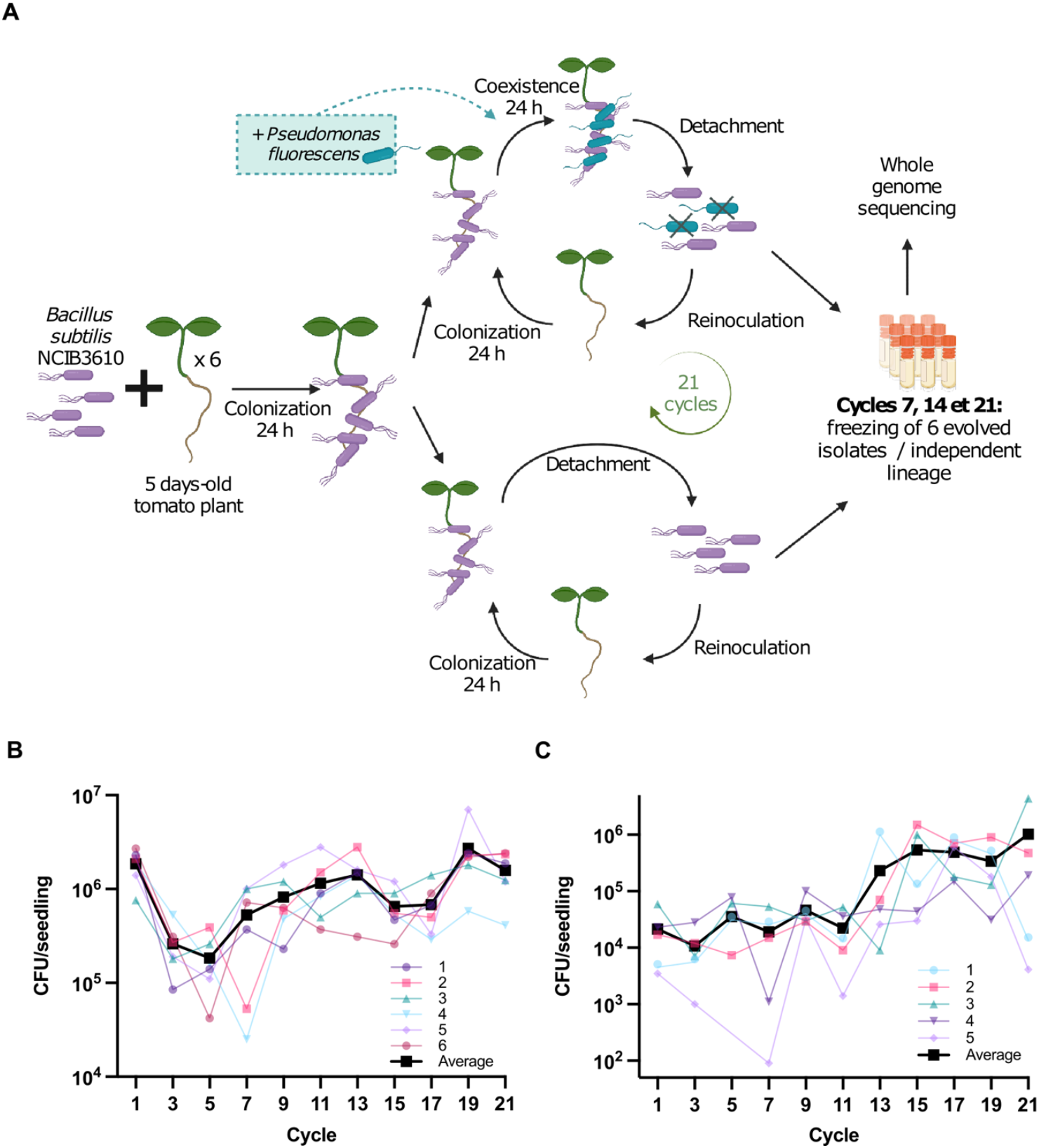
Evolution of *B. subtilis* on tomato roots. (A) Depiction of the directed evolution of *B. subtilis* in presence of *P. fluorescens* WCS365 or alone on tomato plant roots. (See *Methods* for detailed description of experiment). (B) CFU of *B. subtilis* colonizing the root at every two cycles for the experimental evolution alone or (C) with *P. fluorescens*. Fig 2A was created with BioRender.com

As previously observed by Blake et al. 2021 (31), we noticed that different morphological variants emerged during the evolution and rapidly dominated the population with the ancestral strain morphology disappearing only a few cycles after the first appearance of a new variant. After 7, 14, and 21 strain cycles, six evolved isolates per lineage were cryopreserved by selecting differently shaped colonies after plating. The morphology of the frozen isolates was then compared, and we selected at least one strain of each morphology present in a lineage at a specific cycle to be sequenced. A total of 78 evolved isolates (36 for evolution alone and 42 for paired evolution) were sequenced. The analysis revealed that 31 of these isolates were clones of at least one other, and 12 isolates were perfect match of the ancestral strain. Of note, the ancestral strain was also re-sequenced to establish a strong basis for the analysis. Single nucleotide polymorphisms (SNPs) were the most frequent type of mutation for both evolution experiments, representing 69.4% of all mutations, compared with 3.2%, 3.2%, and 24.2% for insertions, synonymous variants, and deletions respectively. Overall, 62 different mutations were identified, of which 39 were found in the BRPE, 19 with the BRE, and 3 were shared between both (Tables S1 and S2). Even though more mutations were acquired by isolates evolved with *P. fluorescens*, the ratios of SNPs over other different types of mutations between both evolution experiments were very similar (70% vs 69%).

Our analysis revealed that the BRPE led to more isolates with 2 or more mutations (17 out of 27) than the BRE (6 out of 13). Interestingly, all lineages from BRE bared a conserved SNP that appeared before the 7^th^ cycle, contrasting with the paired evolution, where only two lineages maintained a conserved mutation from cycle 7 to 21 (Tables S1 and S2). All those observations suggest that the adaptation in a tri-partite system, with the root and *Pseudomonas* (BRPE), took longer than the adaptation to the host alone (BRE).

### Mutations in biofilm regulation genes are conserved amongst evolved isolates

By cycle 21, we observed that SNPs in the negative biofilm regulator genes *sinR* or *ywcC* were found in all lineages. For the BRE, conserved SNPs in *ywcC* appeared in 3 independent lineages and SNPs in *sinR* appeared in the 3 other lineages (Table S1). In the BRPE, SNPs in the two genes were also found in 3 lineages each but were not mutually exclusive since one lineage contained SNPs in both genes (Tables 2 and S2). Apparition of these SNPs in parallel evolutions and lineages suggest that they confer an advantage to root colonization. We chose to further examine one evolved isolate per lineage for most lineages (BRE; Table 1) and three pairs of evolved isolates, each coming from different lineage and which display increase SNPs throughout the cycles (BRPE; Table 2). The latter allowed us to examine the effect of single SNPs versus a combination.

**Table 1.**
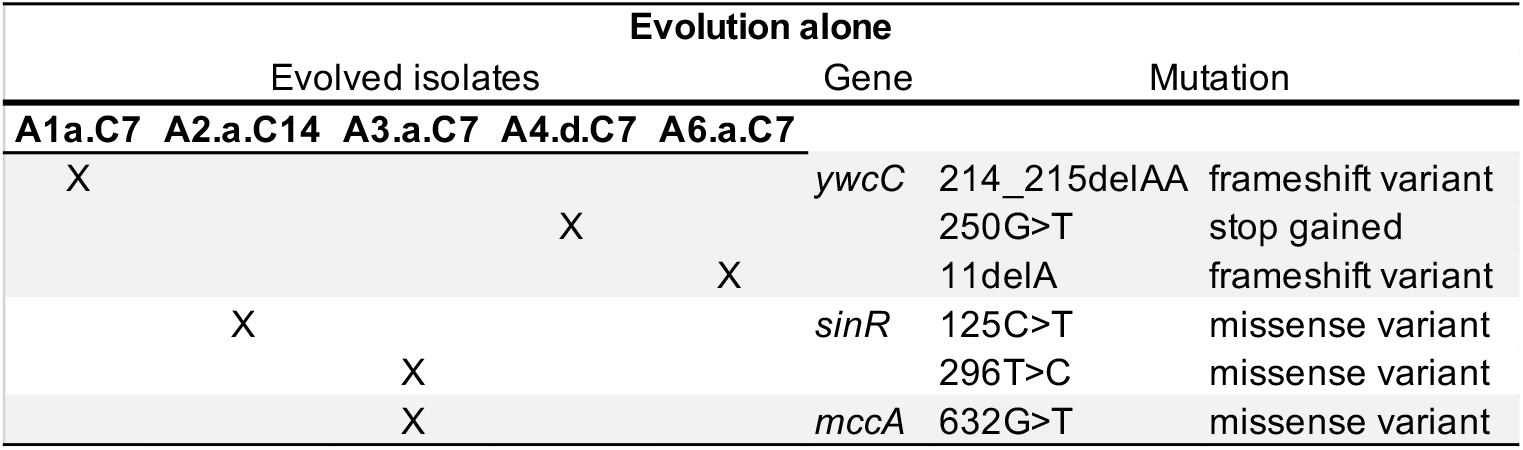
SNPs identified in the genomes of isolates evolved alone (BRE) on tomato roots.

**Table 2.** SNPs identified in the genomes of isolates evolved with *P. fluorescens* (BRPE) on tomato roots.

| Evolution with *P. fluorescens* |  |  |  |  |  | Mutation |
| --- | --- | --- | --- | --- | --- | --- |
| Evolved isolates |  |  |  |  | Gene |  |
| P3.a.C7 | P3.a.C14 | P4.a.C7 | P4.b.C14 | P5.a.C14 | P5.d.C21 |  |
| X | X |  | X |  |  | *ywcC* 11delA frameshift variant |
|  | X |  |  |  |  | *sinR* 296T>C missense variant |
|  |  |  |  | X | X | 170C>T missense variant |
|  |  |  |  |  | X | *dxs* 1246G>T missense variant |
|  | X |  |  |  |  | *dxr* 386C>T missense variant |
|  |  | X | X |  |  | *pksR* 5041T>C missense variant |

### Evolved isolates show enhanced colonization in a plant-independent manner

The competitivity of the evolved isolates was first tested on tomato plants with or without *P. fluorescens*, in an experimental set up similar to the directed evolution assay. As shown in Fig 3A, most evolved isolates showed a 2-fold increase in root colonization capacity when inoculated alone. Similar results were obtained for colonization assays in presence of *P. fluorescens*, even from isolates that were evolved without the selective pressure of the competitor (Fig 3B). This observation suggests that the capacity to compete with *P. fluorescens* strongly correlates with an enhanced root biofilm formation capacity.

**Fig 3.**
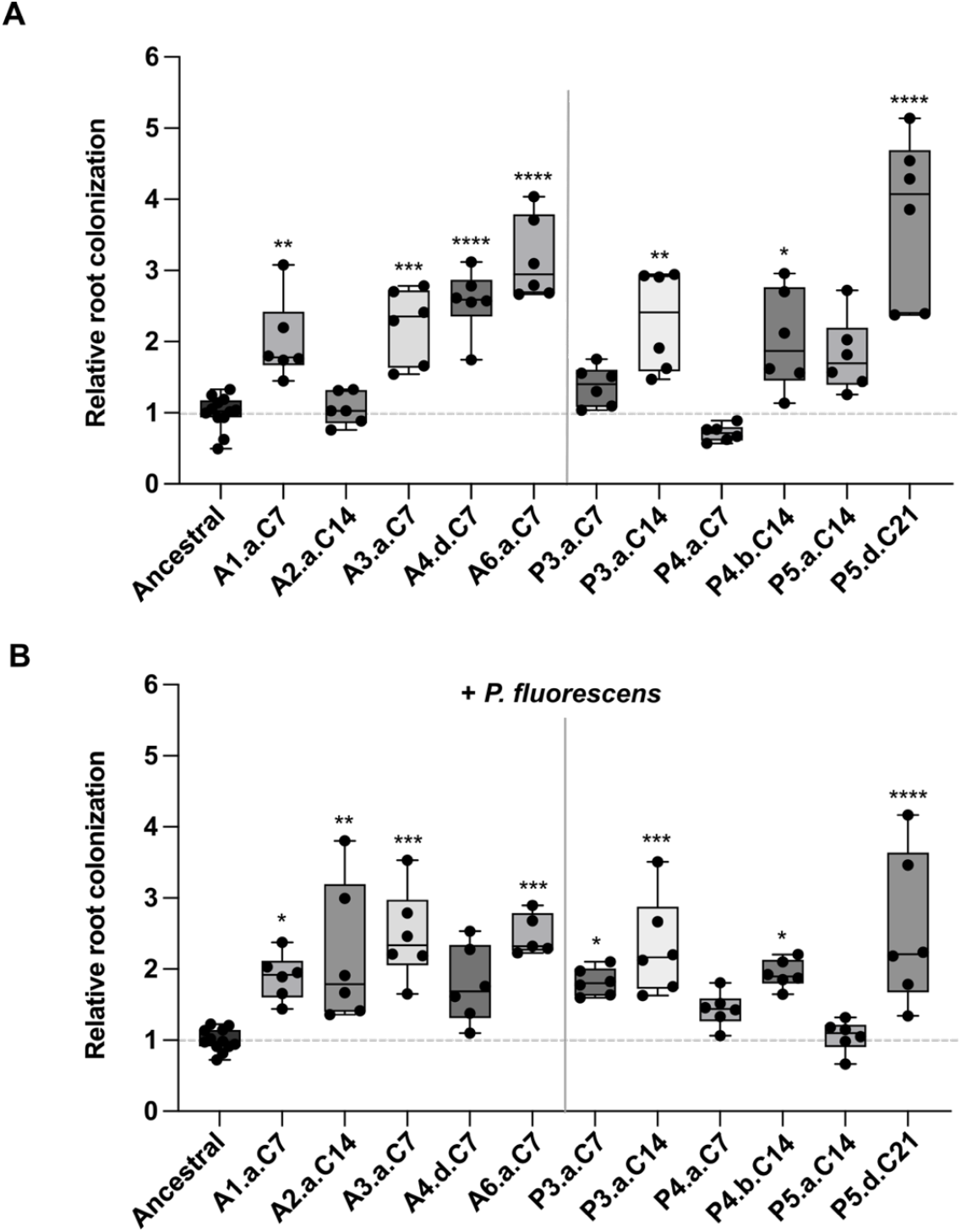
Tomato root colonization by evolved isolates. Tomato root colonization by evolved isolates from the evolution alone (starts with A) and the evolution with *Pseudomonas* (starts with P) without (A) or with *P. fluorescens* WCS365 (B). 5-6 days-old tomato plants were inoculated with ancestral strain or evolved isolates constitutively expressing mKATE2 and incubated for 24 h in a growth chamber. Bacteria were detached and quantified by fluorescence intensity for each plant. Relative root colonization was calculated by dividing the fluorescence intensity of each plant on the average intensity for the ancestral strain of the same experiment. Asterisks indicate statistically significant differences compared to ancestral strain, * = P < 0.05, ** = P < 0.01, *** = P < 0.001, **** = P < 0.0001, ANOVA followed by Dunnett’s post hoc test.

Hu et al. 2023 (32) recently reported that *B. subtilis* adapts differently to various plant species, as some mutations were detected more frequently when the evolution was performed on *A. thaliana* versus tomato plant roots. Another recent study observed that isolates evolved on *A. thaliana* roots did not have an improve root colonization of tomato plants (31). Here, as shown in Fig 4, our tomato-evolved isolates from both evolution appeared to have some increased capacity to colonize *A. thaliana* roots. The results are however more variable, which might be due to the strength of the phenotype. Curiously, only two evolved isolates showed significant improvement of *A. thaliana* root colonization when inoculated alone (Fig 4A), but many showed increase colonization compared to the ancestral strain when *P. fluorescens* was present (Fig 4B). These observations suggest that evolution on tomato roots does not frequently lead to increase colonization of *A. thaliana*, but the increased competitivity in presence of *P. fluorescens* is conserved on both plants.

**Fig 4.**
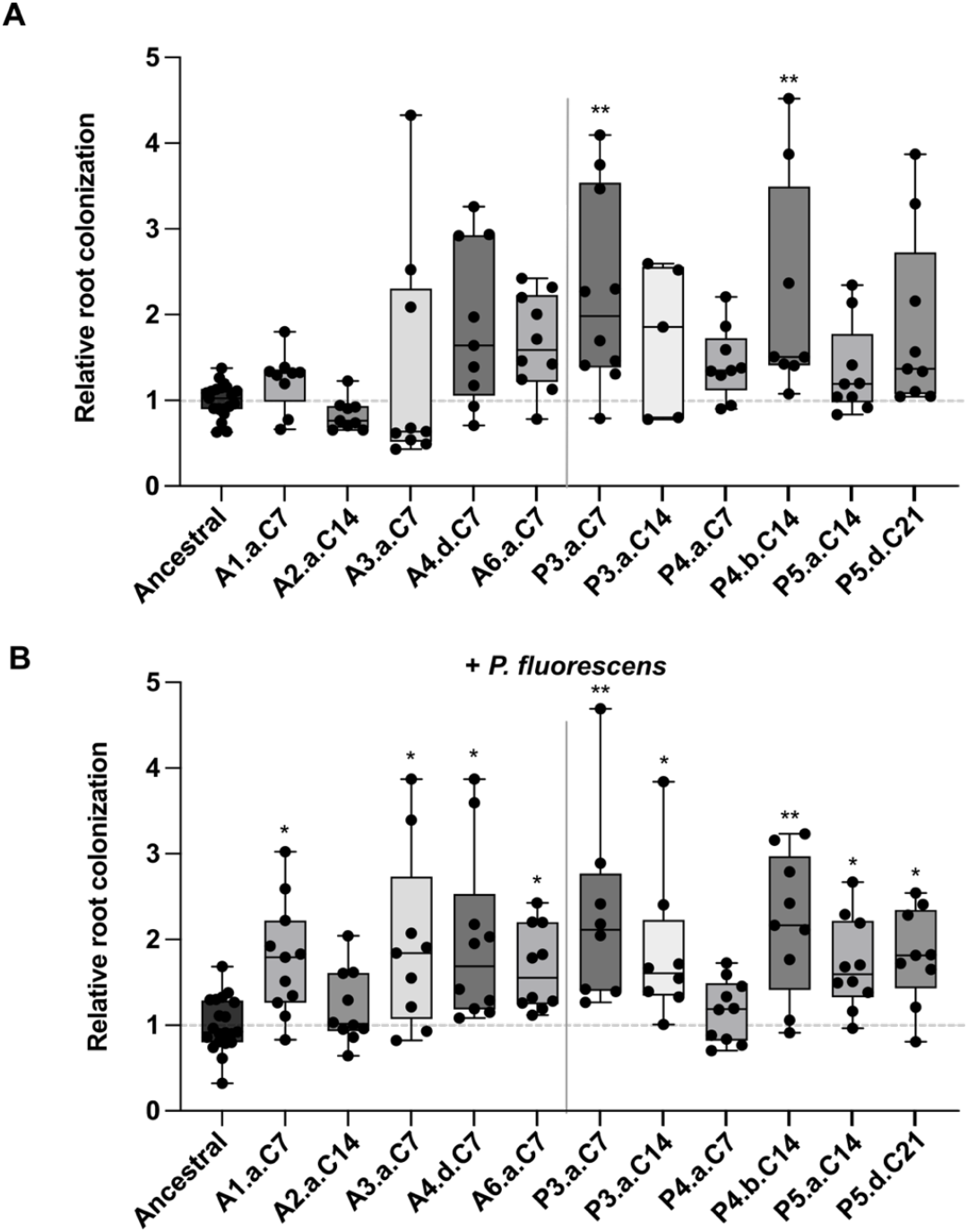
*A. thaliana* root colonization by evolved isolates. *A. thaliana* root colonization by evolved isolates from the BRE (starts with A) and the BRPE (starts with P), without (top panel; A) or with *P. fluorescens* WCS365 (bottom panel; B). 6 days-old plants were inoculated with ancestral strain or evolved isolates constitutively expressing mKATE2 (with or without *P. fluorescens*) and incubated for 24 h in a growth chamber. Whole root pictures were taken using Zeiss Axio Observer Z1 microscope, root area was delimited, and relative root colonization was calculated by dividing the mean fluorescence intensity by pixel of each root on the average intensity for the ancestral strain of the same experiment. Asterisks indicate statistically significant differences compared to ancestral strain, * = P < 0.05, ** = P < 0.01, Kruskal-Wallis followed by Dunn’s test.

### Evolved isolates compete better with other *Pseudomonas* species on roots

As shown in Fig 1, other *Pseudomonas* species also negatively affect *Bacillus* root colonization. Thus, three of the evolved isolates showing the strongest colonization in presence of *P. fluorescens* (P3.a.C14, P4.b.C14 and P5.d.C21) were also tested in presence of four other *Pseudomonas*, inoculated individually at the same time. The species used, *P. protegens* and *P. stutzeri*, have with *P. fluorescens* WCS365 large difference in gene content, and thus likely use different antagonistic strategies against *B. subtilis* (45). Despite this, as shown in Fig 5 we observed that our evolved isolates showed stronger root colonization than the ancestral strain in presence of either of these *Pseudomonas* strain. This important result hints that the adaptation to *P. fluorescens* led to a general mechanism that also confers advantages in root colonization when another antagonistic member of the same genus is present.

**Fig 5.**
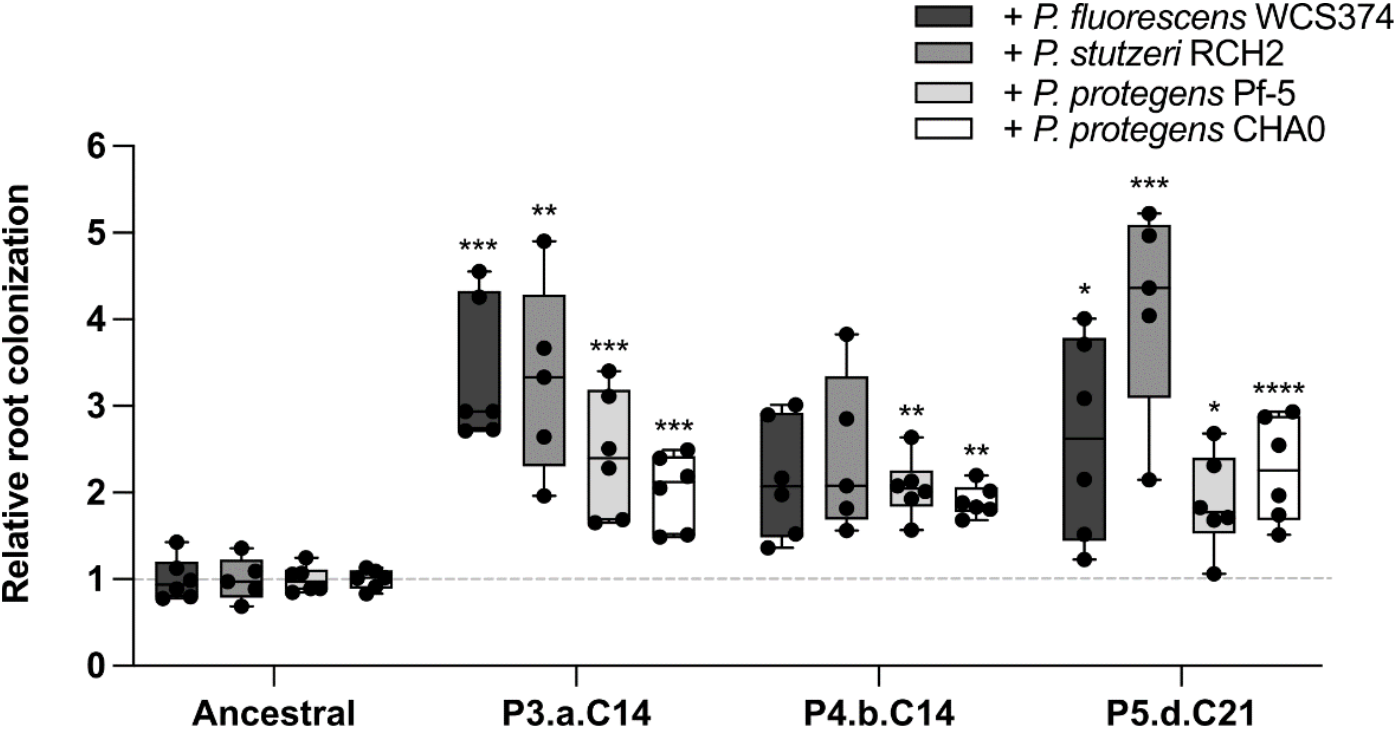
Mutations in biofilm regulators confer fitness toward various *Pseudomonas* spp. Root colonization in presence of other *Pseudomonas* spp. Tomato plants were inoculated as previously described, with mKATE2 expressing ancestral strain or evolved isolates, and four *Pseudomonas* species. * = P < 0.05, ** = P < 0.01, *** = P < 0.001, **** = P < 0.0001, ANOVA followed by Dunnett’s post hoc test (except in 5A for + *P. fluorescens* WCS374 and + *P. stutzeri* RCH2: Kruskal-Wallis followed by Dunn’s test).

### Impacts of the SNPs on the biofilm regulators functionality

Evolved isolates showing better colonization in presence of *Pseudomonas* carry SNPs in *ywcC, sinR* and *dxr* (P3.a.C14), *ywcC* and *pksR* (P4.b.C14) and *sinR* and *dxs* (P5.d.C21; see Table 2). For *ywcC*, all the SNPs resulted in a frameshift or a stop codon, thus suggesting a loss-of-function, while the SNPs present in the other genes were missense variants. Since these SNPs provided an advantage when colonizing roots alone and in the presence of different *Pseudomonas*, we examined deletion mutants for *ywcC, sinR, pksR* and recreated the double mutants as found in the evolved isolates (Fig 6). Interestingly, only Δ*ywcC* and Δ*ywcC ΔpksR* strains showed increased root colonization compared to the wild-type strain, further supporting that the SNPs in *ywcC* resulted in a loss-of-function. Conversely, a *sinR* deletion mutant did not confer an increase in root colonization and even negated the advantage provided by *ΔywcC*, suggesting that the SNPs in *sinR* only partially reduced its function. Accordingly, the morphologies of Δ*sinR* strains did not recapitulate the morphologies of the evolved isolates with *sinR* SNPs, further supporting this hypothesis (Fig S1).

**Fig 6.**
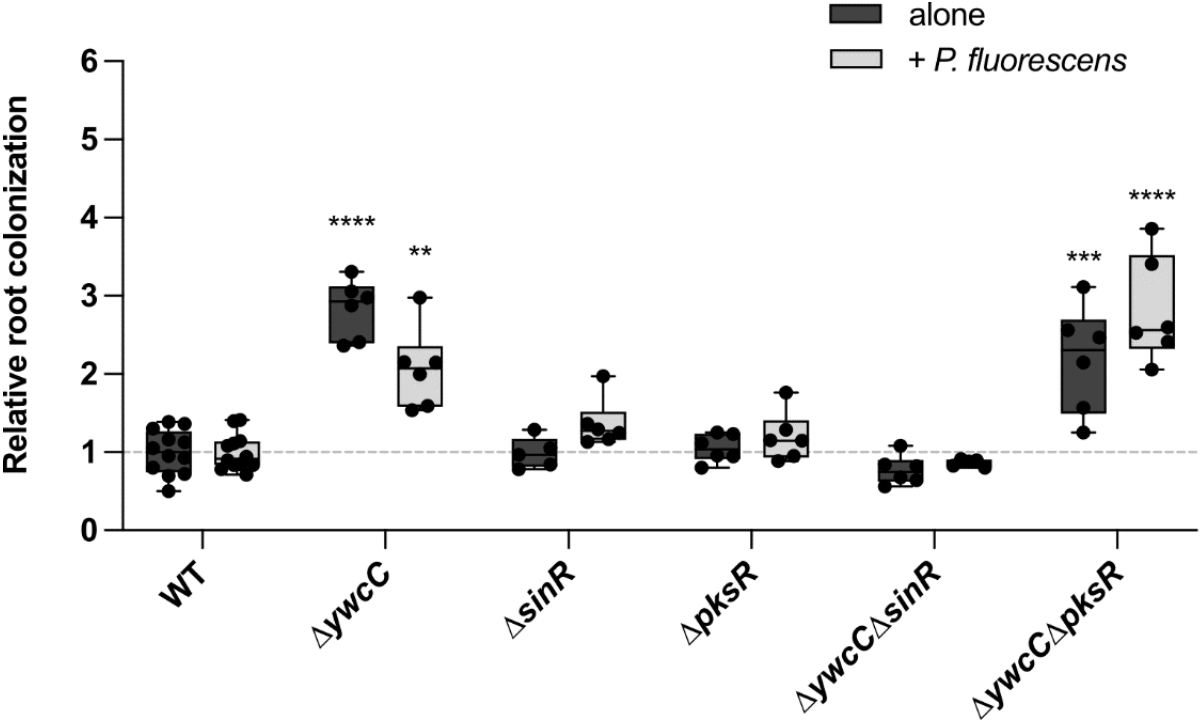
Deletion of biofilm regulator gene *ywcC* improve tomato root colonization. Relative colonization of tomato roots by deletion mutants of genes of interest in the study, done as descrided earlier. Relative root colonization was calculated by dividing the fluorescence intensity of each plant on the average intensity for the ancestral strain of the same experiment. * = P < 0.05, ** = P < 0.01, *** = P < 0.001, **** = P < 0.0001, ANOVA followed by Dunnett’s post hoc test

## Discussion

In this study, we showed that the antagonism often observed between *Bacillus* spp. and *Pseudomonas* spp. is also observed on plant roots since two different *Pseudomonas* species negatively impacted *Bacillus* root colonization of both *A. thaliana* and tomato (Fig 1). This result could reflect direct antagonism via antimicrobial mechanisms, similar to what was previously described using pairwise interaction (21,22,36). The weaker inhibitory effect of *P. fluorescens* might be due to its lower number of secondary metabolites (46,47). However, the antagonism could also be indirect; these *Pseudomonas* could be better adapted to the root microenvironment, or capable of establishing faster on the roots, causing a physical displacement of *B. subtilis*.

The directed evolution on tomato roots, performed as a monoculture or in presence of *P. fluorescens* WCS365, allowed the emergence of different morphotypes of *B. subtilis* through time. Seven out of the total eleven lineages obtained at cycle 21 had only one morphotype present, while three other lineages had two morphotypes, and one lineage had four morphotypes (Tables S1 and S2). Whole genome sequencing revealed that multiple independent lineages from both evolutions shared SNPs in the same few genes, suggesting that evolutionary pressure came from the host and the experimental conditions more than the presence of the competitor. A few of our isolates evolved on tomato also had increased *A. thaliana* root colonization, showing the possibility of one strain being more fit on two different plants. Of note, presence of 0.05% glycerol in our evolution assay might have relieved the pressure to adapt to specific carbon compounds present in the exudates.

Either *ywcC* or *sinR* were affected by SNPs in all independent lineage from both evolution experiments, stressing their importance for root colonization adaptation. Both genes encode for negative regulators of biofilm genes, and their deletion result in hyper-robust biofilm formation (24,48,49). The YwcC protein is a TetR-type transcriptional repressor that inhibits *slrA*, a *sinI* paralog. When expressed, SlrA inhibits SinR thereby derepressing matrix genes (48). Consequently, it was proposed that YwcC responds to a yet unknown environmental signal and alleviate its repression, allowing SlrA to transiently induce matrix production for the whole cell population (48,50). YwcC also repress poly-y-glutamic acids production, which were shown to play a notable role in plant root colonization (26). Evolved isolates harboring a frameshift mutation in the *ywcC* gene demonstrated enhanced root colonization compared to the ancestral strain, and their morphology suggested robust biofilm formation (Fig S1). Thus, the mutations acquired during the evolution likely result in a loss of function, which is further supported by the fact that Δ*ywcC* recapitulated *ywcC*\* evolved isolates phenotypes.

On the contrary, *sinR* loss-of-function did not increased root colonization, even in presence of *P. fluorescens* or in combination with Δ*ywcC*, pointing toward a milder impact of the SNPs on SinR function. Certain evolved isolates with *sinR* SNP, such as A2.a.C14 from the evolution alone (125C>T; P42L in the DNA binding domain; (51)), had no impact on root colonization while displaying a different morphology than the ancestral strain. However, one specific mutation, found in both evolution assays, seemed to enhance root colonization (296T>C; L99S). This mutation might impair SinR capacity to form a functional tetramer and thus, partially reduce its capacity to repress target genes (52). These observations are in accordance with the observation similar results of evolved isolates on *A. thaliana* roots in which robust biofilm formers had an increase in root colonization (30).

The paired evolution led to the emergence in lineage 4 of a conserved mutation in the *pksR* gene. This gene is part of the *pks* gene cluster, encoding for an enzymatic megacomplex that synthesizes bacillaene, an antimicrobial compound (53). However, this SNP alone (evolved isolate P4.a.C7) did not confer any advantage when colonizing tomato or *A. thaliana* roots, contrary to when this mutation was combined with a SNP *ywcC* (P4.b.C14), suggesting that the latter SNP was responsible of the adaptation.

Finally, SNP in two essential genes involved in the MEP pathway of isoprenoid biosynthesis were identified in two independent isolates of the paired evolution showing stronger roots colonization (P3.a.C14 and P5.d.C21) (Table 2, *dxr* and *dxs*). While the increased fitness by P3.a.C14 can be attributed to a frameshift mutation in *ywcC*, in P5.d.C21 the *dxs* mutation is accompanied only by a *sinR* SNP that has little to no effect by itself (see P5.a.C14 in Fig3B and Fig4B), indicating this *dxs* SNP had positive effect on the roots. Since *dxs* and *dxr* are essential genes, the mutations are at best a partial loss-of-function; further experiments will be required to determine the reason behind these beneficial effects.

Overall, directed evolution performed alone or in presence of *P. fluorescens* lead to parallel evolution of SNPs in a few genes that conferred a mild deregulation of biofilm genes. These evolved isolates showed increase presence on the root compared to the ancestral strain, alone or with various *Pseudomonas* species. Our study indicates that rhizosphere fitness in presence of a competitor is closely linked to robust root colonization and physical establishment.

## Materials and Methods

### Bacterial strains and culture conditions

Strains used in this study are listed in Supplementary Table 3. *B. subtilis* strains were routinely cultivated in lysogeny broth (Luria-Bertani LB; 1% w/v tryptone, 0.5% w/v yeasts extract, 0.5% w/v NaCl) at 37 °C in agitation for 3 h. *Pseudomonas* strains were cultivated in LB at 30 °C in agitation for 3 h. Experimental evolution and all root colonization assays were performed in half strength Murashige and Skoog (MS) basal medium (2.22 g L^−1^, Sigma) supplemented with 0.05% glycerol. When necessary, antibiotics were used at the following concentrations: spectinomycin (100 μg·mL^−1^), kanamycin (10 μg·mL^−1^), chloramphenicol (5μg·mL^−1^), erythromycin (1μg·mL^-1^).

#### Seedling preparation

*A. thaliana* ecotype Col-0 and tomato (*Solanum lycopersicum*, Rutgers, McKenzie seeds, CA and Greta’s Family Gardens, CA) were used in this study. Seeds were surfaced sterilized with 70% v/v ethanol followed by 0.3% v/v sodium hypochlorite then rinsed three times with sterile MiliQ water and plated on Murashige and Skoog basal medium (4.44 g L^−1^, Sigma) 0.8% w/v agar supplemented with 0.05% w/v glucose. Plates were incubated in a growth chamber with a day/night cycle of 12 h light at 25°C and 12 h dark at 20°C. *A. thaliana* seedlings for 7 days and tomato seedlings for 5 or 6 days.

#### Strains constructions

All genetically modified strains were made by transferring genetic constructions into NCIB3610 or evolved isolates using SPP1-mediated generalized transduction (54). All deletion mutants used in this study were purchased from the Bacillus Genetic Stock Center (BGSC) collection (http://www.bgsc.org) in the *B. subtilis* 168 background. Pseudomonas strains *P. fluorescens* WCS374, *P. stutzeri* RCH2 and *P. protegens* CHA0 were obtained from Cara Haney (UBC).

#### Experimental evolution

Tomato seedlings were transferred to 6-well plates containing 4 mL of 1/2 MS + 0.05% v/v glycerol (2 seedlings/plate in opposite corners to prevent contamination between lineages). *B. subtilis* NCIB3610 culture was washed in PBS and inoculated at a final optical density at 600 nm (OD_600_) of 0.02. Plates were put on a shaker at 90 rpm in the growth chamber in the same conditions than for seedlings preparation. **Evolution alone (BRE):** After 24 h, the root was detached from the stem using tweezers, rinsed in PBS and sonicated at 30% amplitude for 30 s in a microtube containing 1 mL PBS. 133 μL of the bacterial suspension was inoculated on a new plant and the suspension was diluted and plated to follow the number of bacteria transferred every cycle. **Evolution with Pseudomonas (BRPE):** After 24 h, *P. fluorescens* WCS365 was inoculated in each well containing a *B. subtilis* colonized root at a final OD_600_ of 0,0002, which correspond approximately to a 100 : 1 Bacillus : Pseudomonas ratio, and reincubated in the growth chamber for 24 h. Any lower Bacillus : Pseudomonas ratio lead to the rapid disappearance of the *Bacillus* population, suggesting an evolutionary pressure (not shown). Root was then detached, washed, and sonicated (40% amplitude, 60 s), which killed 99% of the *Pseudomonas*. 133 μL of the bacterial suspension was then inoculated on a new plant and the suspension was diluted and plated to follow the number of bacteria transferred every cycle. For the cycles 3 to 21, *P. fluorescens* was added after 24 h at each cycle at a final OD_600_ of 0,00002, or approximately 7,3 × 10^3^ CFU/mL (OD_600_ of 0,6 = 2,2 × 10^3^ CFU/mL). Considering the *B. subtilis* CFU number on the root (see Fig 2c), this *P. fluorescens* inoculum corresponds to a Bacillus : Pseudomonas ratio of around 1:10 to 100:1, depending on the lineage and the cycle.

### Whole genome sequencing

Genomic DNA was extracted from 9 mL of 3 h cultures using Monarch® Genomic DNA Purification Kit. Paired-end (evolution alone) or single-end (evolution with *Pseudomonas*) libraries were prepared using the NEBNext® Ultra™ II DNA Library Prep with Sample Purification Beads. Fragment reads were generated on an Illumina NextSeq sequencer using TG NextSeq® 500/550 High Output Kit v2 (300 cycles) (RNOmique Platform, UdeS). The methodology for processing the reads drew inspiration from the GenPipes framework (55). We first implemented a quality-based trim of the reads using fastp 0.21.0 (56), setting the parameters to --cut_right --cut_window_size 4 --cut_mean_quality 30 --length_required 30. The trimmed reads were subsequently mapped to the reference *Bacillus subtilis* 3610 and its plasmid pBS32, procured from GenBank (CP020102.1 and CP020103.1), using BWA-MEM v0.7.10 (57) with no modification of the default parameters. Upon alignment, we sorted the reads with Picard v1.123, available online at https://broadinstitute.github.io/picard/, and realignment was executed with the GATK v3.7 RealignerTargetCreator and IndelRealignerSingle tools (58). Following the identification and marking of duplicate reads with Picard v1.123, we utilized GATK v3.7 for haplotype identification. Subsequently, their impacts were evaluated using SnpEff v4.1 (59).

#### Colonization assays

Evolved isolates formed very robust biofilm on root surface, so CFU counting to quantify root colonization was unreliable since clumps of bacteria could not be separated by sonication. We developed two methods that allowed us to quantify relative root colonization regardless of clumps, using a constitutive fluorescent reporter that was transduced into all strains tested. **For tomato:** *B. subtilis* and evolved mutants were transduced with the *amyE*::P_*hyperspank*_-*mkate2*, and then inoculated on roots as described in the evolution assay on tomato plants. For the condition in presence of *Pseudomonas*, both species were inoculated at the same time and at the same OD (OD_600_= 0.02). After 24 h, the roots were detached and transferred in a microtube containing 500 μL PBS with three 3 mm glass beads. Roots were vortexed for 30 s and sonicated at 40% amplitude for 20 s. 200 μL of the supernatant was transferred to a 96-well plate and the fluorescent intensity was measured with a TECAN Spark monochromator-based with an excitation wavelength of 590 nm and emission wavelength of 638 nm, bandwidth 20. A calibration curve using this method is shown in Fig S2. **For *A. thaliana*:** The assays were performed in 48-well plates containing 280-290 μL of media. Bacteria were either detached from root using sonication (amplitude 30%, 10 pulses 1-s pulse and 1-s rest time), diluted, and plated on solid LB media to allow CFU counting, or bacterial colonization was quantified in microscopy (see Microscopy section).

### Microscopy

To visualize bacteria on root surface for Fig 1, seedlings were examined with a Zeiss Axio Observer Z1 microscope equipped with a 20X/0.8 Plan-Apochromat objective, and whole root pictures were taken with a Zeiss Axiocam 506 mono. The fluorescence signal was detected using a Cy3 filter (ex: 545/25, em: 605/70). All images were taken at the same exposure time, processed identically, and prepared for presentation using ImageJ. Each image is representative of at least 9 roots colonization assays performed in three independent experiments. Quantification was done using Fiji. Root area was delimited, and relative root colonization was calculated by dividing the mean fluorescence intensity on the average intensity for the ancestral strain of the same experiment. Fig S1 was obtained using the stereo microscope Leica M165 FC with the LeicaMC170HD camera.

#### Statistical analysis

Statistical analyses were performed using GraphPad Prism 9. Comparisons were done as specified in the figure legends. Normality was tested using Shapiro-Wilk test.

## Acknowledgements

We thank members of Beauregard laboratory and Rodrigue laboratory for helpful discussions. We also thank Daniel Garneau for technical advice and processing of images. We thank Julie Beaudin for critical reading of the manuscript. This work was supported by NSERC discovery grant RGPIN-2020-07057 to P.B.B.

