## Supplementary Tables and Figures for "Adaptative Laboratory Evolution reveals biofilm regulating genes as key players in *B. subtilis* root colonization"

**Supplementary material**

Table S1. Mutations identified in the genomes of isolates from the BRE on tomato roots.

| Gene | Mutation | | Lineage 1 | | | Lineage 2 | | | | | | | Lineage 3 | | | | Lineage 4 | | | | Lineage 5 | | | Lineage 6 | | |
|---|---|---|---|---|---|---|---|---|---|---|---|---|---|---|---|---|---|---|---|---|---|---|---|---|---|---|
| | | | 1.a.C7 | 1.a.C14 | 1.a.C21 | 2.a.C7 | 2.a.C14 | 2.b.C14 | 2.c.C14 | 2.e.C14 | 2.a.C21 | 2.b.C21 | 3.a.C7 | 3.a.C14 | 3.b.C14 | 3.a.C21 | 4.a.C7 | 4.d.C7 | 4.a.C14 | 4.a.C21 | 5.a.C7 | 5.a.C14 | 5.a.C21 | 6.a.C7 | 6.a.C14 | 6.a.C21 |
| ipk | c.241G>T | missense_variant | | | | | | | | | | | | X | | | | | | | | | | | | |
| B4U62_22510 | c.523_*31del | stop_lost&inframe_deletion&splice_region_variant | | | | | | | | | | | | X | | | | | | | | | | | | |
| srfAC | c.3386C>A | missense_variant | | | | | | | | | | | | | | | | | | | | | | | | |
| yloN | c.157C>T | missense_variant | | | | | | X | | | | | | | | | | | | | | | | | | |
| codY-flgB | n.1691261C>T | intergenic_region | | | | | | | | | | | | | | | | | | | | | | | | |
| yodI | c.104G>C | missense_variant | | | | | | | X | | | | | | | | | | | | | | | | | |
| sinI-sinR | n.2552667G>A | intergenic_region | | | | | | | | | | | | | | | | | | | | | | | | |
| sinR | c.29G>T | missense_variant | | | | | | | | | | | | | | | | | | | X | X | X | | | |
| | c.125C>T | missense_variant | | | | | X | | X | | | X | | | | | | | | | | | | | | |
| | c.296T>C | missense_variant | | | | | | | | | | | X | X | X | X | | | | | | | | | | |
| | c.311G>T | missense_variant | | | | X | | X | | X | X | | | | | | | | | | | | | | | |
| mccA | c.632G>T | missense_variant | | | | | | | | | | | X | | | | | | | | | | | | | |
| sdpA | c.77delA | frameshift_variant | | | | | | | | | | | | | | | | | | | | | | | | |
| epsC | c.1289T>G | missense_variant | | | | | | | | | | | | | | | | | | | | | | | | |
| ptkA | c.448G>T | stop_gained | | | | | | | | X | | | | | | | | | | | | | | | | |
| ywcC | c.250G>T | stop_gained | | | | | | | | | | | | | | | | X | X | | | | | | | |
| | c.214_215delAA | frameshift_variant | X | X | X | | | | | | | | | | | | | | | | | | | | | |
| | c.215delA | frameshift_variant | | | | | | | | | | | | | | | | | | X | | | | | | |
| | c.11delA | frameshift_variant | | | | | | | | | | | | | | | | | | | | | | X | X | X |
| aldX | c.923_967delTCGAAAGAGGGGCGGAAGTGGTGTTCGGCGGCGTATTCGATGCCA | disruptive_inframe_deletion | | | | | | | | | | | | | | | | | | X | | | | | | |

### Table S2. Mutations identified in the genomes of isolates from the BRPE on tomato root.

| Gene | Mutation | | Lineage 1 | | | | | Lineage 2 | | | | | | | | | Lineage 3 | | | | | Lineage 4 | | | | | | Lineage 5 | | | | |
|---|---|---|---|---|---|---|---|---|---|---|---|---|---|---|---|---|---|---|---|---|---|---|---|---|---|---|---|---|---|---|---|---|
| | | | 1.a.C7 | 1.a.C14 | 1.c.C14 | 1.a.C21 | 1.b.C21 | 2.e.C7 | 2.a.C14 | 2.b.C14 | 2.c.C14 | 2.d.C14 | 2.d.C14 | 2.e.C14 | 2.f.C14 | 2.a.C21 | 3.a.C7 | 3.b.C7 | 3.a.C14 | 3.e.C14 | 3.a.C21 | 4.a.C7 | 4.b.C14 | 4.a.C21 | 4.b.C21 | 4.e.C21 | 4.f.C21 | 5.a.C7 | 5.a.C14 | 5.b.C14 | 5.a.C21 | 5.d.C21 |
| ftsH | c.1269C>T | synonymous_variant | | | | | | | | | | | | X | | | | | | | | | | | | | | | | | | |
| cshA | c.1357G>T | missense_variant | | | | | | | | X | | | | | | | | | | | | | | | | | | | | | | |
| yesZ | c.1444G>T | stop_gained | | | | | | | | | X | | | | | | | | | | | | | | | | | | | | | |
| slp-ykzW | n.1534070C>A | intergenic_region | | | | | | | | | | | | | | | | | | | | | | X | | | | | | | | |
| ylzA | c.53_90delGGATGATTTCGATTGTCAGCCCGGAGTCTGCGCCAATC | frameshift_variant | | | X | | | | | | | | | | | | | | | | | | | | | | | | | | | |
| flhA | c.926C>G | stop_gained | | | | | | X | | | | | | | | | | | | | | | | | | | | | | | | |
| dxr | c.386C>T | missense_variant | | | | | | | | | | | | | | | | | X | | X | | | | | | | | | | | |
| pksM | c.1804C>T | missense_variant | | | | | | | | | | | | | | | | | | X | | | | | | | | | | | | |
| pksR | c.5041T>C | missense_variant | | | | | | | | | | | | | | | | | | | | X | X | X | X | X | X | | | | | |
| ppsC | c.3511G>A | missense_variant | | | | | | | | | | | | | | X | | | | | | | | | | | | | | | | |
| ppsB | c.4737G>T | synonymous_variant | | | | | | | | | | | | | | X | | | | | | | | | | | | | | | | |
| ppsA-dacC | n.1998147delT | intergenic_region | | | | | | | | | | X | | | | | | | | | | | | | | | | | | | | |
| yotL | c.107G>T | missense_variant | | X | | | | | | | | | | | | | | | | | | | | | | | | | | | | |
| dxs | c.1246G>T | missense_variant | | | | | | | | | | | | | | | | | | | | | | | | | | | | | | X |
| | c.857C>G | missense_variant | | | | | | | | | | | | | | | | | | | | | | | | | | | | | X | |
| sinI-sinR | n.2552661G>T | intergenic_region | | | | | | | X | X | X | X | X | X | X | X | | | | | | | | | | | | | | | | |
| sinR | c.127T>G | missense_variant | | X | X | | X | | | | | | | | | | | | | | | | | | | | | | | | | |
| | c.170C>T | missense_variant | | | | | | | | | | | | | | | | | | | | | | | | | | | X | | X | X |
| | c.296T>C | missense_variant | | | | | | | | | | | | | | | | | X | X | X | | | | | | | | | | | |
| bscR-yrhH | n.2777799delT | intergenic_region | | | | | | | | | | | | | | | | | | | | | | X | | | | | | | | |
| dhbE | c.776C>A | missense_variant | | | | X | | | | | | | | | | | | | | | | | | | | | | | | | | |
| dhbC | c.695G>T | missense_variant | X | | | | | | | | | | | | | | | | | | | | | | | | | | | | | |
| cssR | c.351G>C | synonymous_variant | | | | | | | | | | | | | | | | X | | | | | | | | | | | | | | |
| epsD | c.467_488delTTCAAAGGCCGGGCGGAAGGAC | frameshift_variant | | | | | | | | | | | | | X | | | | | | | | | | | | | | | | | |
| epsC | c.141_142delTG | frameshift_variant | | | | | | | | | | | X | | | | | | | | | | | | | | | | | | | |
| degU | c.436G>T | stop_gained | | | | | | | | | | | | | | | | | | | | | | | X | | | | | | | |
| tagO | c.747A>G | missense_variant | | | | | | | | | | | | | | | | | | | | | | | | | X | | | | | |
| gtaB | c.347G>T | missense_variant | | | | | | | | | | | | X | | | | | | | | | | | | | | | | | | |
| | c.351_352delAC | frameshift_variant | | | | | | | | | | | | X | | | | | | | | | | | | | | | | | | |
| | c.354_355insCT | frameshift_variant | | | | | | | | | | | | X | | | | | | | | | | | | | | | | | | |
| | c.382G>T | missense_variant | | | | | | | | | | | | X | | | | | | | | | | | | | | | | | | |
| | c.384A>T | synonymous_variant | | | | | | | | | | | | X | | | | | | | | | | | | | | | | | | |
| | c.393T>A | synonymous_variant | | | | | | | | | | | | X | | | | | | | | | | | | | | | | | | |
| B4U62_1924[illegible] | c.26T>G | missense_variant | | | | | | | | | | | | X | | | | | | | | | | | | | | | | | | |
| B4U62_1924[illegible] | c.48T>C | synonymous_variant | | | | | | | | | | | | X | | | | | | | | | | | | | | | | | | |
| ywcC | c.215delA | frameshift_variant | | | | | | | | | | | | | | | | | | | | | | | | | | | | X | | |
| | c.11delA | frameshift_variant | | | | | | | | | | | | | | | X | | X | X | X | | X | X | X | X | X | | | | | |
| yxiO | c.685A>T | missense_variant | | | | | | | | | | | | | | | | | | | | | | | | | | | | | X | |

Table S3. Strains used in this study.

| Strain | Genotype | Reference |
|---|---|---|
| NCIB 3610 | WT/undomesticated | Lab stock |
| PB389 | 3610 *amyE*::$P_{hyperspank}$-mKATE2 | Lab stock |
| PB127 | 3610 *ywcC*::kan | YC295[1] |
| MP57 | 3610 *ywcC*::kan *amyE*::$P_{hyperspank}$-mKATE2 | This study |
| PB18 | 3610 *sinR*::spc | Lab stock |
| MP63 | 3610 *sinR*::spc *amyE*::$P_{hyperspank}$-mKATE2 | This study |
| BKK17220 | 168 *pksR*::kan | BGSC[2] |
| MP65 | 3610 *pksR*::kan | This study |
| MP66 | 3610 *pksR*::kan *amyE*::$P_{hyperspank}$-mKATE2 | This study |
| BKE17220 | 168 pksR::erm | BGSC[2] |
| MP67 | 3610 *ywcC*::kan *pksR*::erm *amyE*::$P_{hyperspank}$-mKATE2 | This study |
| MP60 | 3610 *ywcC*::kan sinR::spc | This study |
| MP68 | 3610 *ywcC::*kan *sinR*::spc *amyE*::$P_{hyperspank}$-mKATE2 | This study |
| *Pseudomonas fluorescens* | WCS365 | Lab stock |
| *Pseudomonas fluorescens* | WCS374 | Cara Haney |
| *Pseudomonas capeferrum* | WCS358 | Lab stock |
| *Pseudomonas protegens* | Pf-5 | Lab stock |
| *Pseudomonas protegens* | CHA0 | Cara Haney |
| *Pseudomonas stutzeri* | RCH2 | Cara Haney |

Antibiotics resistance abbreviations: Kanamycin (kan), spectinomycin (spc), erythromycin (erm)

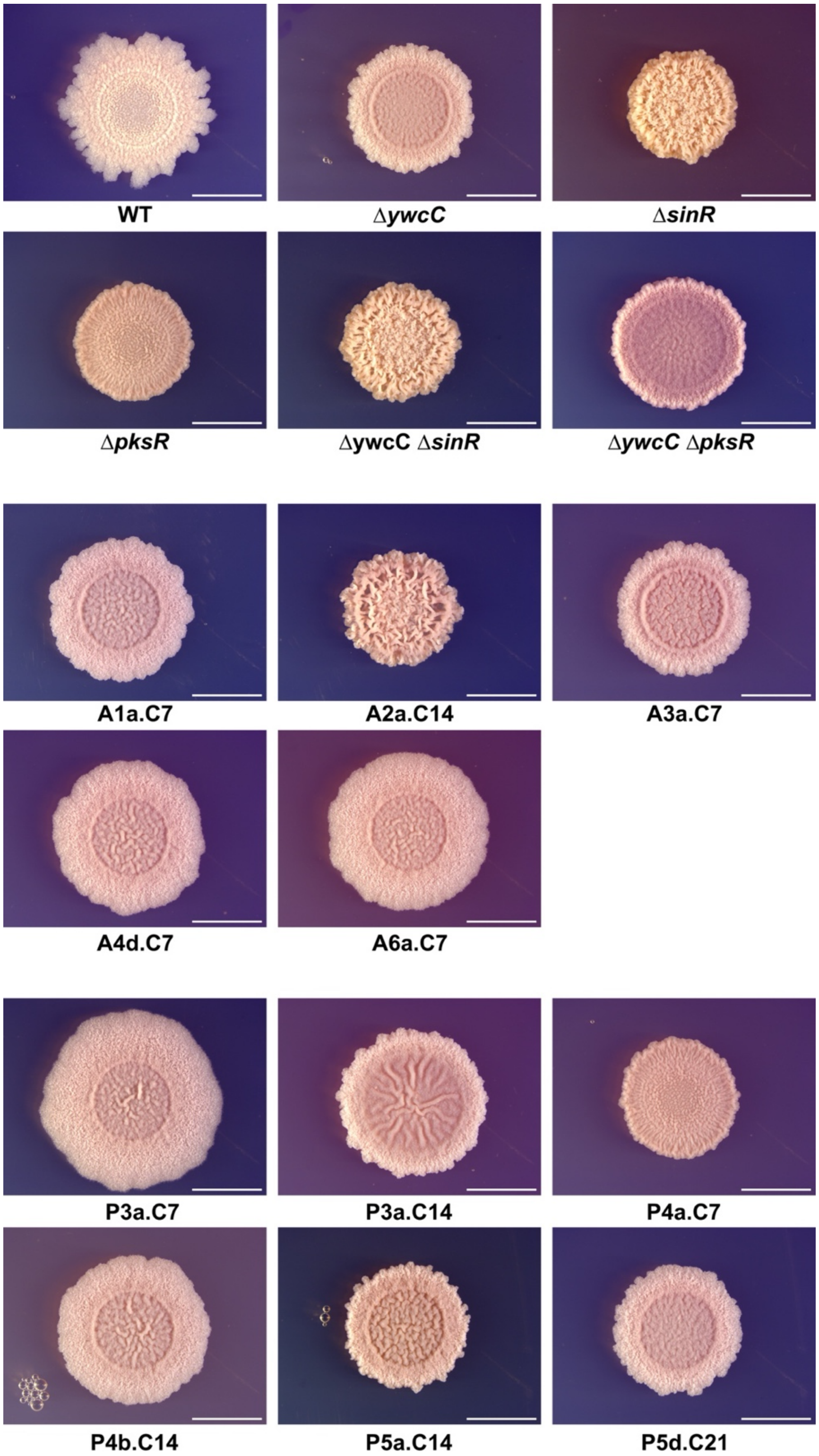


**Fig S1. Morphotypes of deletion mutants and evolved isolates**

Representative pictures of colony morphotypes of deletion mutants or evolved isolates grown on MS agar + 0.5% glutamate + 0.5% glycerol for 48 h. Scale bar is 5 mm for all images.

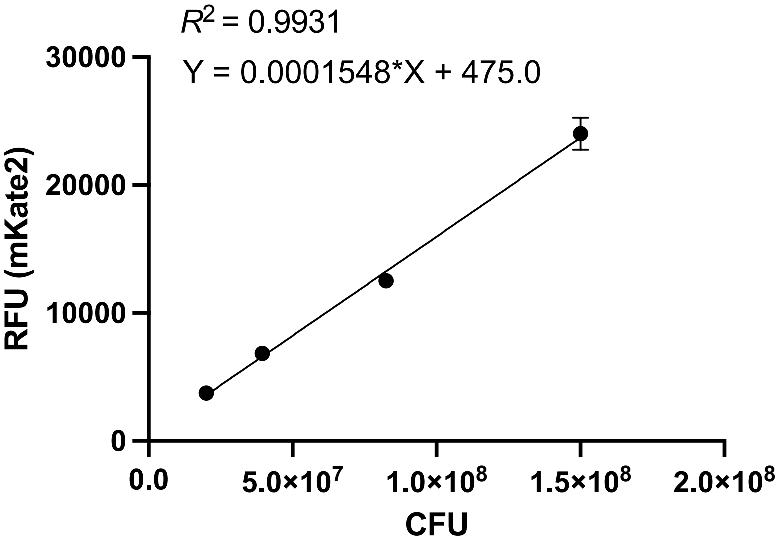


**Fig S2. Relative fluorescence unit (RFU) is a quantitative tool which correlates with bacterial counts.** RFU was measured using TECAN Spark monochromator-based (plate reader) at emission: 590 nm, excitation: 638 nm, and bacterial counts was followed by dilution and plating. Representative experiment of 3 replicates is presented.
